# L-GIREMI uncovers RNA editing sites in long-read RNA-seq

**DOI:** 10.1101/2022.03.23.485515

**Authors:** Zhiheng Liu, Giovanni Quinones-Valdez, Ting Fu, Mudra Choudhury, Fairlie Reese, Ali Mortazavi, Xinshu Xiao

## Abstract

Using third-generation sequencers, long-read RNA-seq is increasingly applied in transcriptomic studies given its major advantage in characterizing full-length transcripts. A number of methods have been developed to analyze this new type of data for transcript isoforms and their abundance. Another application, which is significantly under-explored, is to identify and analyze single nucleotide variants (SNVs) in the RNA. Identification of SNVs, such as genetic mutations or RNA editing sites, is fundamental to many biomedical questions. In long-read RNA-seq, SNV analysis presents significant challenges, due to the well-known relatively high error rates of the third-generation sequencers. Here, we present the first study to detect and analyze RNA editing sites in long-read RNA-seq. Our new method, L-GIREMI, effectively handles sequencing errors and biases in the reads, and uses a model-based approach to score RNA editing sites. Applied to PacBio long-read RNA-seq data, L-GIREMI affords a high accuracy in RNA editing identification. In addition, the unique advantage of long reads allowed us to uncover novel insights about RNA editing occurrences in single molecules and double-stranded RNA (dsRNA) structures. L-GIREMI provides a valuable means to study RNA nucleotide variants in long-read RNA-seq.

## Introduction

Adenosine-to-Inosine (A-to-I) RNA editing is one of the most common RNA modification types in human cells, which greatly diversifies the transcriptome [1]. A-to-I RNA editing is catalyzed by enzymes encoded by the adenosine deaminase acting on RNA (*ADAR*) gene family in Metazoans [2–5]. ADAR proteins recognize and bind to double-strand RNAs (dsRNAs) to deaminate adenosines into inosines [6–9]. Most RNA editing sites in human cells occur in *Alu* repeats [10–13], the most abundant type of short interspersed elements (SINEs).

A-to-I RNA editing occurs in both coding and non-coding regions, with diverse functional roles in human cells [1,14,15]. The impact of recoding sites (i.e., those that alter protein-coding sequences) has been relatively well studied, many of which alter protein function [16]. It is now known that RNA editing in non-coding regions can influence gene expression, such as by affecting alternative splicing [17–19] or RNA stability [20–22]. In addition, RNA editing affects microRNA maturation, leading to the crosstalk between RNA editing and RNA interference [23,24]. Recently, regulation of the immunogenicity of dsRNAs is emerging as an important aspect of RNA editing function [25–27]. Given the diverse functions of RNA editing, abnormal editing patterns have been reported for numerous diseases, such as neurological diseases, autoimmune disorders, and cancers [1,14,28–30].

Next-generation sequencing technologies, especially RNA-sequencing (RNA-seq), have greatly facilitated the discovery of RNA editing events [31–34]. To date, more than 16 million RNA editing events have been cataloged in human transcriptomes [35]. In order to segregate RNA editing sites from single-nucleotide polymorphisms (SNPs) in the genome, many previous methods required sequencing of both DNA and RNA of a sample. In our previous work, we developed a method, namely GIREMI, to accurately identify RNA editing events using a single short-read RNA-seq dataset without genome sequencing data of the corresponding sample [36].

With the development of third-generation sequencing (TGS) technologies, long-read RNA-seq methods recently emerged as powerful tools to study RNA biology. Pacific Biosciences (PacBio) and Oxford Nanopore Technologies (ONT) are two main representatives of the TGS platforms. Different from short-read RNA-seq methods, long-read RNA-seq interrogates full-length transcripts without breaking the RNAs into small fragments, thus preserving transcript structures [37]. As a result, long-read RNA-seq overcomes the transcript assembly ambiguities inherent to short-read RNA-seq, greatly improving the understanding of transcriptome diversity [38].

A number of methods have been developed to analyze long-read RNA-seq data, primarily focusing on transcript isoform identification and their abundance analysis [39–42]. Another application, which is significantly under-explored, is to identify and analyze single nucleotide variants (SNVs) in the RNA. Identification of SNVs, such as genetic mutations or RNA editing sites, is fundamental to many biomedical questions. In long-read RNA-seq, SNV analysis presents significant challenges, due to the well-known high error rates of the third-generation sequencers.

We present L-GIREMI (Long-read GIREMI), the first method to identify RNA editing sites in long-read RNA-seq (without the need of genome information). L-GIREMI effectively handles sequencing errors and biases in the reads, and uses a model-based approach to score RNA editing sites. L-GIREMI allows investigation of RNA editing patterns of single RNA molecules, co-occurrence of multiple RNA editing events, and detection of allele-specific RNA editing. This method provides new opportunities to study RNA nucleotide variants in long-read RNA-seq.

## Results

### Overview of the L-GIREMI method

Linkage patterns between alternative alleles of RNA variants in the mRNA differ for different types of RNA variants. For example, a pair of SNPs within the mRNA are generally expected to possess perfect allelic linkage. In contrast, non-genetic RNA variants, such as RNA editing sites, do not generally show significant allelic linkage with each other nor with SNPs (unless allele-specific editing exists). In our previous work, we showed that these properties can be employed to distinguish RNA editing sites from genetic variants using short-read RNA-seq data [43]. Long-read RNA-seq affords a major advantage in capturing such allelic linkage since multiple variants in the same mRNA can be covered by each read. Leveraging this feature of long-read RNA-seq, we developed the L-GIREMI method to identify RNA editing events using this data type.

After a typical read mapping procedure (e.g., via minimap2 [44]), the L-GIREMI algorithm is mainly composed of four steps (Fig. 1). First, the strand of each read was examined and corrected if necessary (see Methods). Second, mismatch sites in the BAM file were obtained and pre-filtered according to common practices in detecting RNA editing sites using RNA-seq data [45,46]. In the third step, the mutual information (MI) between pairs of mismatch sites in the same gene was calculated. Specifically, an average MI was calculated for each unknown mismatch relative to putative SNPs (from dbSNP) covered by the same reads. Similarly, MI of pairs of putative heterozygous SNPs (from dbSNP) was also obtained. Since most RNA editing events are expected to occur independently of the allelic origin of the mRNA, the above MI for an RNA editing site should be smaller than that between two heterozygous SNPs. Thus, the two types of MI values were compared to predict RNA editing sites among the unknown mismatches. The predicted RNA editing sites then served as training data for the fourth step, where a generalized linear model (GLM) was derived. Sequence features and allelic ratios of candidate sites were included as predictive variables in the GLM and a score was calculated for each mismatch (Methods).

**Figure 1.**
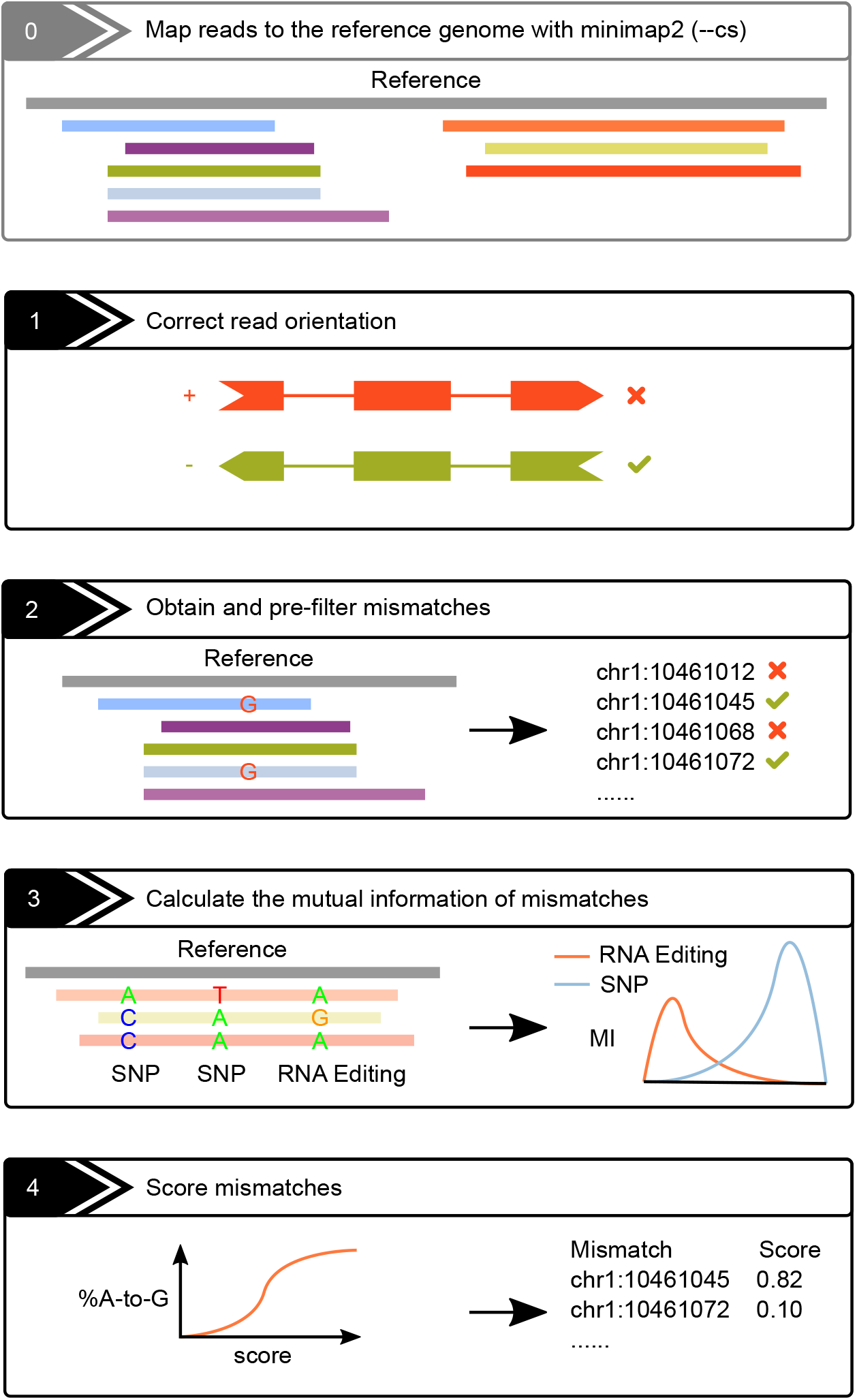
The schematics of the L-GIREMI algorithm (see text for details).

### Performance evaluation of L-GIREMI

We first tested the performance of L-GIREMI using a dataset derived from the brain sample of an Alzheimer’s disease (AD) patient ((PacBio Sequel II, data available at PacBio, 4,277,293 reads). As expected [47], the majority of reads harbored at least one mismatch or indel (Fig. S1a). On average, 14 mismatches, 38 deletions, and 11 insertions were found in each read (Fig. S1b). Thus, the nature of long-read RNA-seq presents substantial challenges in resolving *bona fide* nucleotide variants.

L-GIREMI overcame these challenges and effectively detected RNA editing sites from the dataset. As shown in Fig. 2a, upon the initial screen of nucleotide variants, all 12 types of single nucleotide mismatches were detected in the mapped reads, with the A-to-G type (likely due to A-to-I editing) constituting only a small fraction. The L-GIREMI built-in filters (Methods) were applied to remove sites possibly arising from sequencing errors (Step 2, Fig. 1), which greatly improved the %A-to-G among all mismatches (Fig. S2a). Subsequently, MI values were calculated for mismatch sites that shared at least 6 reads with putative heterozygous SNPs (defined as dbSNPs with allelic ratio between 0.35 and 0.65 in the data). As a comparison, the MI values of pairs of putative heterozygous SNPs were also calculated. As shown in Fig. 2b, the MI distribution of unknown mismatches was well separated from that of putative heterozygous SNPs. Thus, using the MI distribution of SNPs, we calculated an empirical p value for each mismatch site and identified those with p<0.05 as candidate RNA editing sites. These sites were in turn used as training data for the GLM model (Methods). In total, 28,584 RNA editing sites were detected in the AD dataset, with 98.1% being A-to-G mismatches (Fig. 2c). The high fraction of A-to-G sites among all predicted editing sites attests to the high accuracy of our method. Interestingly, we observed that the %A-to-G sites among all predicted sites increased monotonically with the GLM score (Fig. 2d). By default, L-GIREMI chooses a score cutoff (vertical line in Fig. 2d) for each dataset to optimize the F1 value (Methods). As an alternative approach, a user-defined GLM score cutoff can be provided to achieve a desired %A-to-G, based on the GLM score vs. %A-to-G relationship.

**Figure 2.**
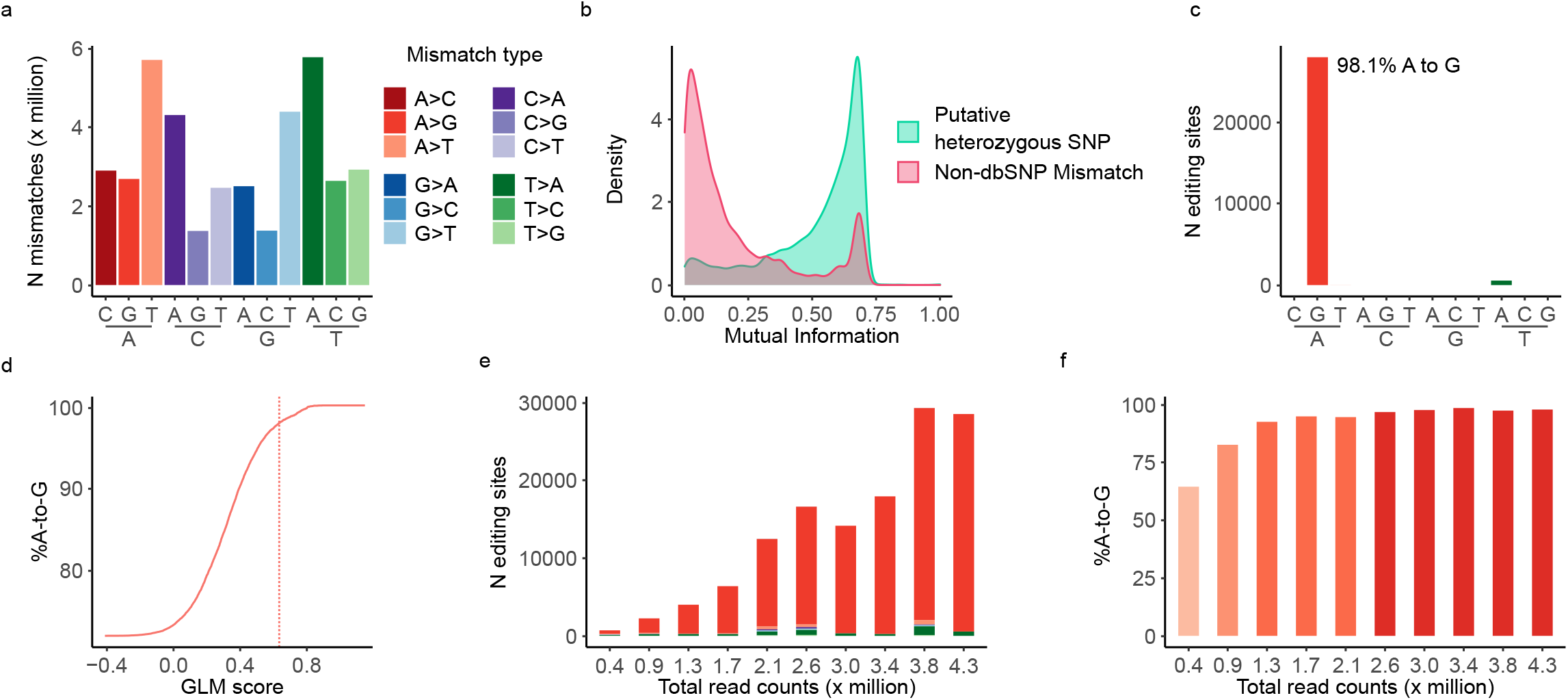
Identification of RNA editing sites in long-read RNA-seq data of the brain sample of an Alzheimer’s disease (AD) patient. (**a**) Raw mismatches detected in the dataset. (**b**) Mutual information for pairs of putative heterozygous SNPs (based on dbSNP) or non-dbSNP mismatches relative to putative SNPs. (**c**) RNA editing sites identified by L-GIREMI. (**d**) %A-to-G among all predicted editing sites vs. GLM score. Dotted line denotes the score cutoff used for (**c**) (0.64). (**e**) Number of RNA editing sites identified given different read coverages (randomly chosen subsets of the AD dataset). (**f**) %A-to-G among the RNA editing sites identified in the subsets in (**e**).

To evaluate the impact of read coverage on the results, we randomly sub-sampled the AD data set to retain different numbers of total reads. As expected, the number of predicted RNA editing sites decreased given reduced read coverage (Fig. 2e). In contrast, the fraction of A-to-G sites among the predicted RNA editing sites (score cutoff determined by F1 score) remains relatively high except for the very low coverage (e.g., 0.4M reads, Fig. 2f). It should be noted that the fraction of A-to-G among the pre-filtered sites (step 2, Fig. 1) is much lower than that of the final L-GIREMI predictions (Fig. S2b), supporting that the MI and GLM steps enhanced the prediction accuracy. In summary, L-GIREMI affords high accuracy in capturing RNA editing sites in long reads for a wide range of total read coverages.

### Identification of RNA editing sites with data of different PacBio platforms

We analyzed three datasets derived from GM12878 cells by the ENCODE project (ENCODE IDs: ENCFF417VHJ, ENCFF450VAU and ENCFF694DIE, 1,673,768 reads, 2,137,168 reads and 2,538,701 reads respectively) via the PacBio Sequel II platform. The 3 datasets had similar error profiles. However, ENCFF417VHJ, which was built using the MaximaH-reverse transcriptase, showed relatively lower levels of indels and mismatches than the other two datasets (Fig. S3a, b).

The GLM score vs. %A-to-G curves were largely monotonic, although exceptions existed (Fig. S3c). The final predicted RNA editing sites for the ENCFF417VHJ dataset had a high level of A-to-G (99.9%), which is much higher than the 31.4% and 37.8% for ENCFF450VAU and ENCFF694DIE respectively (Fig. S3d) (all of which being higher than the % using only pre-filters, Fig. S3e). Therefore, the chemistry used in generating PacBio RNA-seq libraries can greatly influence RNA editing identification. Note that for the latter two datasets, a user-defined GLM score cutoff could be used to achieve a higher accuracy (%A-to-G) in the predicted editing sites. Importantly, we noted that similar data derived from GM12878 cells but using the earlier Sequel platform yielded lower quality and suboptimal RNA editing identification (Fig. S4). Thus, our data suggest Sequel II is a preferred platform for the purpose of RNA editing studies.

### Co-occurrence of RNA editing sites

Consistent with previous reports based on short-read RNA-seq data, the majority of RNA editing sites detected by L-GIREMI in the GM12878 (henceforth, only ENCFF417VHJ was used) and AD datasets were located in non-coding regions and *Alu* elements (Fig. 3a, b) [31,37,48–50]. A prevailing question is whether multiple editing sites of a gene tend to co-occur in a subset of RNA molecules or if their occurrence is largely independent of each other. This question was challenging to address using short reads. In contrast, the long-read RNA-seq data enable a direct examination of this question.

**Figure 3.**
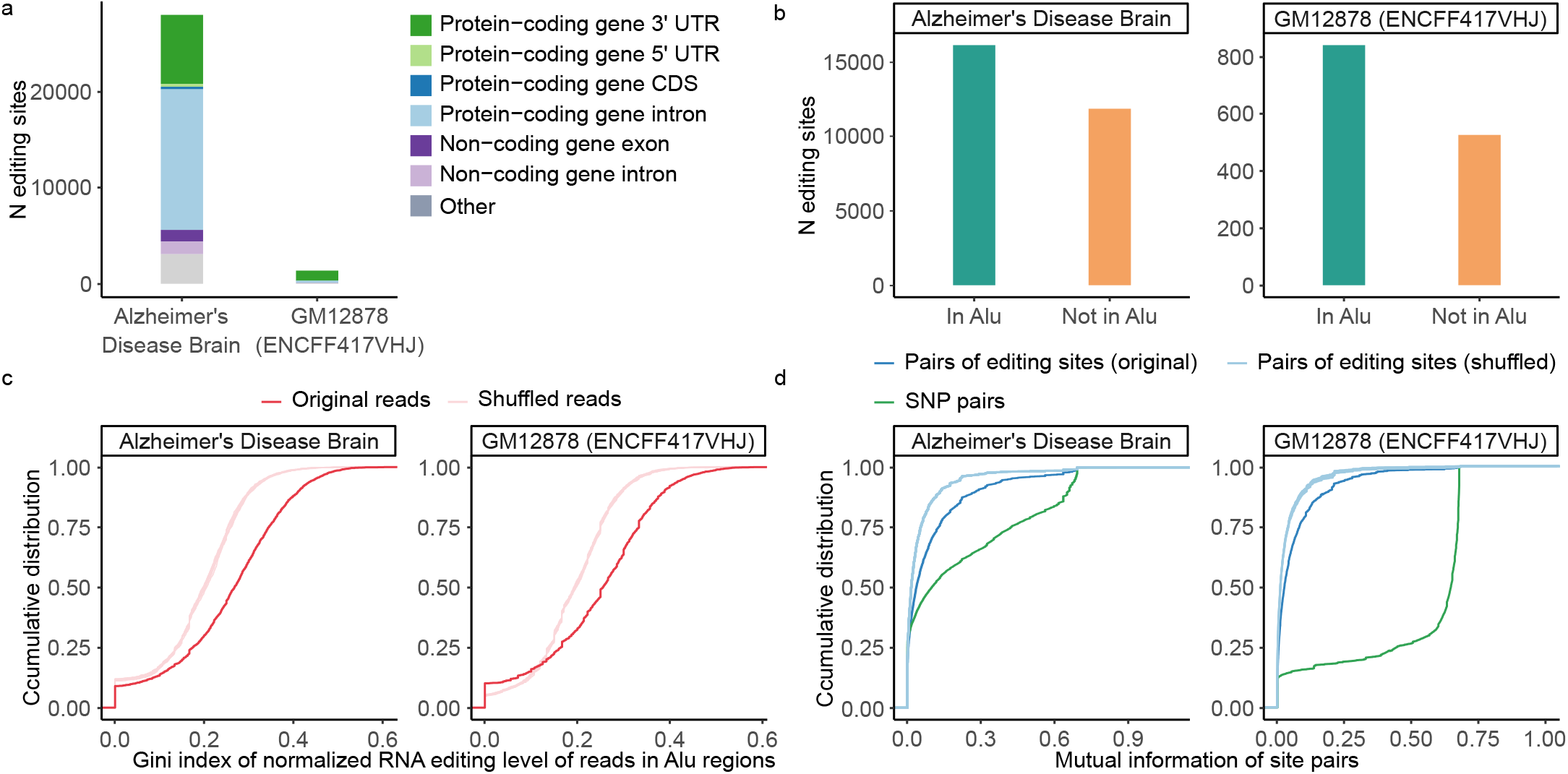
Co-occurence of A-to-I RNA editing sites in *Alu* elements detected by L-GIREMI. (**a**) Genomic context of A-to-I RNA editing sites identified in the AD sample or GM12878 cells. (**b**) Number of RNA editing sites in *Alu* repeats or otherwise. (**c**) Cumulative distribution for the Gini index of *Alu* repeats calculated via read-specific editing ratio. Shuffled data were generated for comparison (Methods), which led to significantly lower Gini index values than the original data (p < 0.001 for both data sets, KS test). (**d**) Cumulative distribution of mutual information of pairs of editing sites or pairs of SNPs in the same gene. Compared to the shuffled controls, both editing sites and SNPs show higher levels of linkage (p < 0.001 for all comparisons, KS test) although the latter were associated with much higher mutual information.

We first asked whether editing sites in the same *Alu* was approximately equally distributed among the reads corresponding to the *Alu*. To this end, we calculated the Gini index for the number of editing sites observed in each read of an *Alu*. Gini index is a measure of inequality among a given set of values (Methods). As background controls, we shuffled the occurrence of the observed editing sites among all reads of the *Alu*. As shown in Fig. 3c, the Gini index of the actual editing profiles is much larger than that of the controls, suggesting the existence of co-occurrence of editing sites in the same *Alu*.

As an alternative method, we examined the mutual information (MI) of pairs of editing sites in a gene and asked whether their MI values are larger than that of randomly shuffled editing sites. Consistent with the above finding (Fig. 3c), the MI values of pairs of editing sites in the same gene are significantly higher than those of the shuffled controls (Fig. 3d). Nonetheless, it should be noted that the MI of editing sites is substantially lower than those between pairs of SNPs (Fig. 3d), which has been established previously [48–52]. This observation still holds if only known editing sites (from REDIportal) were used (Fig. S5). Thus, it is unlikely due to the fact that MI was used in the process of editing site identification. Overall, our results support the existence of co-occurrence of RNA editing sites in the same RNA molecules, to a level significantly higher than random expectations, but significantly lower than genetic linkage.

### Allele-specific RNA editing events detected by L-GIREMI

Long-read RNA-seq allows examination of linkage patterns between any types of RNA variants in the same read. One type of linkage event, allele-specific RNA editing, reflects the existence of genetic determinants of RNA editing, which has been shown in human and mouse tissues [53]. However, it is not clear whether allele-specific RNA editing affects a predominant number of editing sites. We examined this question using the long-read RNA-seq of GM12878 since its whole-genome sequencing data is available. Specifically, we calculated the MI values of all known RNA editing sites in the REDIportal database [35] relative to known SNPs in GM12878 that were detectable in the long-read RNA-seq data. Note that the REDIportal sites were used here instead of editing sites identified in this study so that the source of the editing sites was independent of any linkage calculation. As shown in Fig. 4a, the majority of these REDIportal-defined known editing sites had relatively low MI values, with only a small fraction (∼12%) having MI values greater than 0.3 (the cutoff used in L-GIREMI to identify RNA editing sites in this dataset).

**Figure 4.**
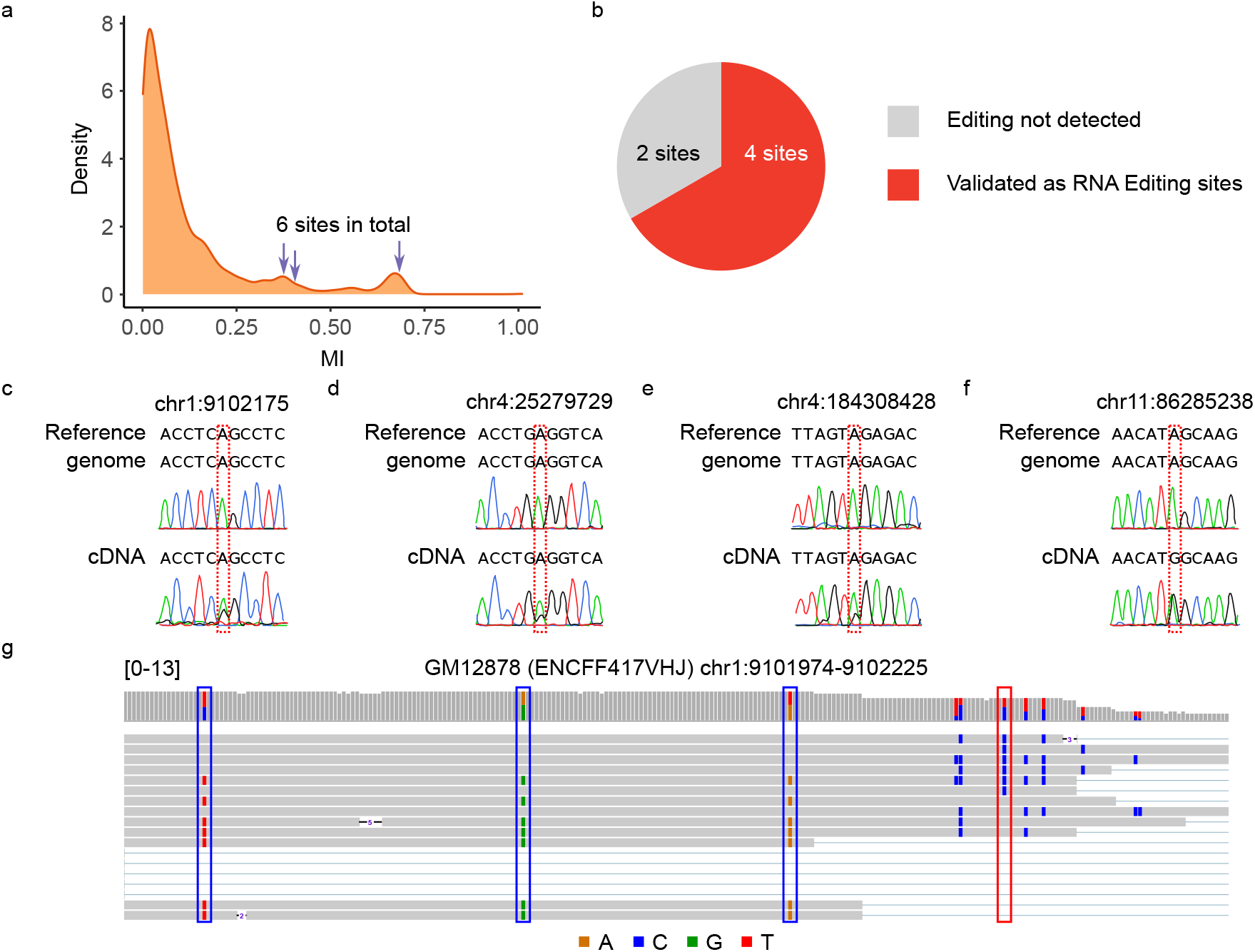
Allele-specific editing reflected in the GM12878 long-read RNA-seq data. (**a**) Mutual information of known editing sites (REDIportal). Six sites with high mutual information were randomly chosen for experimental testing (arrows). (**b**) Summary of experimental validation results. 4 out of 6 sites were confirmed as RNA editing sites. (**c-f**) Sanger sequencing traces for 4 confirmed editing sites. (**d**) IGV plot for an example allele-specific editing event: between the RNA editing site chr1:9102175 (red rectangle) and three heterozygous SNPs (blue rectangle). Note that the gene is on the - strand.

The above results suggest that allele-specific editing may only affect a minority of editing sites. To exclude the possibility that apparent allele-specific RNA editing may be largely due to the existence of genetic variants among REDIportal-defined editing sites (i.e., false positives), we tested 6 likely allele-specific editing sites (arrows, Fig. 4a) using Sanger sequencing. Four of these sites were confirmed as RNA editing sites, whereas two of them were neither edited nor SNPs (thus likely sequencing errors) (Fig. 4b-f). Figure 4g shows the reads harboring an example allele-specific editing event. Therefore, our data suggest that allele-specific RNA editing does exist, although relatively rare.

Since L-GIREMI uses MI as its initial step to predict RNA editing sites, this step excludes allele-specific editing sites. However, such sites may still be captured in the scoring step of L-GIREMI where the GLM model is used for prediction (Fig. S6). In general, L-GIREMI is not recommended for detecting allele-specific editing for novel editing sites. Nonetheless, the MI calculation implemented in L-GIRMI can be used to uncover allele-specific editing of known RNA editing sites.

### dsRNA structures likely affect long read coverage

While inspecting RNA editing sites in the RNA-seq reads, we observed a curious pattern where some long reads showed skipping of a region, often in the vicinity of RNA editing sites. Such skipped regions did not coincide with annotated splicing events. Fig. 5a shows an example in the 3’ UTR of the gene *MREG*. In this example, two *Alu* repeats are located in the skipped region, where many editing sites were identified. This region folds into a strong dsRNA structure (Fig. 5a).

**Figure 5.**
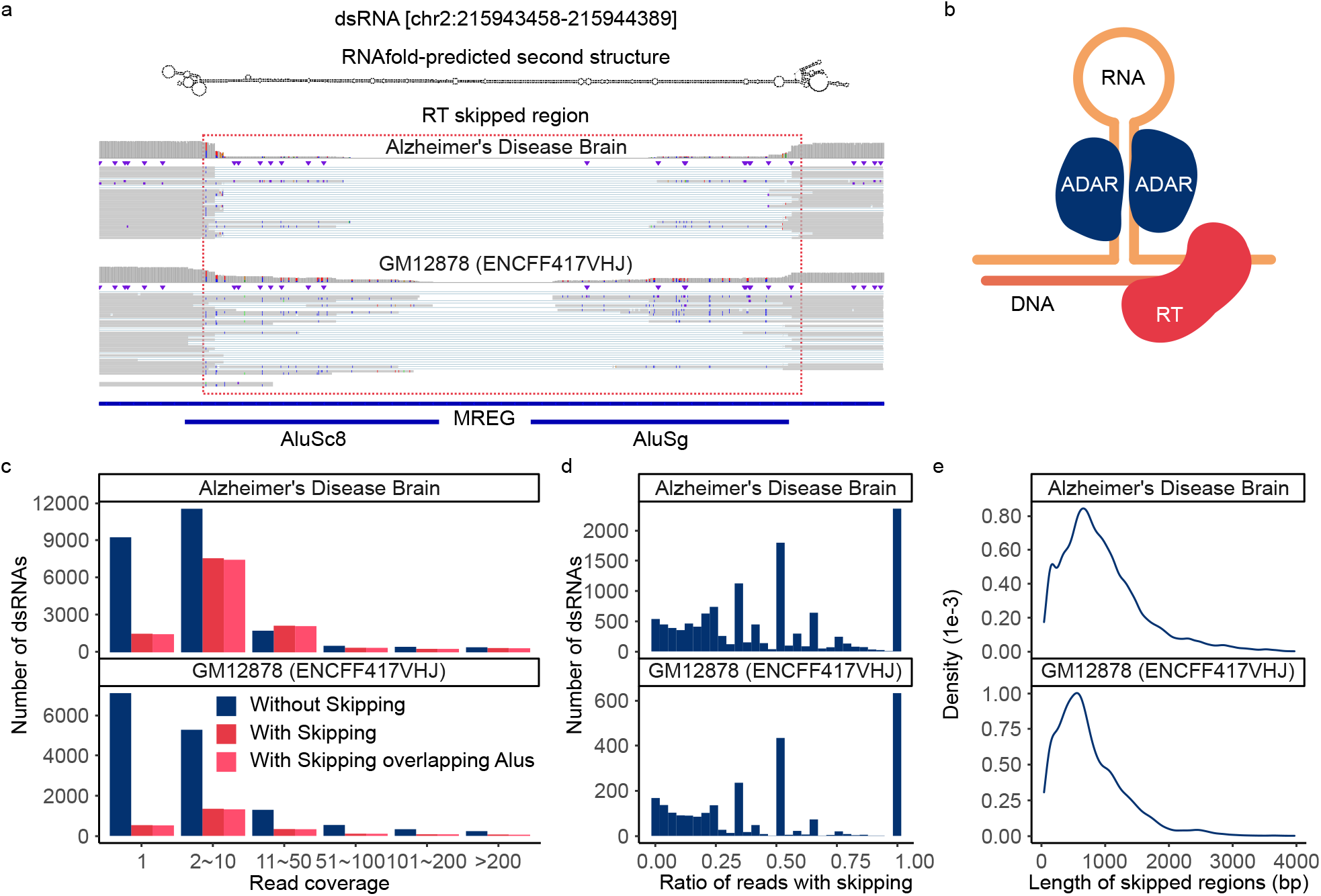
Long-read RNA-seq detected highly structured regions. (**a**) IGV plot for an example where many reads had internal skipping. This region harbors two Alus. The double strand RNA structure predicted by RNAfold is shown. (**b**) Diagram illustrating RT-induced template switching that may induce region-skipping in long reads. (**c**) Read coverage of predicted dsRNAs with or without region-skipping patterns. (**d**) Histogram of dsRNAs with different fractions of reads that showed region-skipping patterns. (**e**) The length of skipped regions within predicted dsRNAs (median = 813, 627 for the AD and GM12878 data, respectively).

We hypothesize that region-skipping in long reads is a consequence of the highly structured nature of the RNA. Indeed, it is known that reverse transcriptase (RT) can generate deletion artifacts in cDNAs, which is caused by intramolecular template switching, an event where RT skips the hairpin structure of the template RNA (illustrated in Fig. 5b) [54]. To further explore this hypothesis, we followed a previously published method to identify dsRNA structures harboring editing-enriched regions [55]. A total of 36,166 and 17,293 predicted dsRNAs were covered by at least 1 read for the AD and GM12878 datasets respectively (Fig. S7). Among these predicted dsRNAs, about 20% overlapped reads with region-skipping (Fig. 5c). In addition, 98.2% and 97.6% of the skipped regions were located between two *Alu* repeats for the AD and GM12878 datasets, respectively (Fig. 5c). For 34.4% and 31.6% of the dsRNAs in AD and GM12878 datasets, respectively, the skipping pattern occurred in ≥50% of their reads (Fig. 5d). The median length of the skipped region was about 600 to 800 basepairs (bp) (Fig. 5e), which is approximately the length of two adjacent *Alu* repeats [11].

## Discussion

Long-read RNA-seq is a powerful means for transcriptome profiling [56–60]. A number of methods have been developed to discover and quantify full-length isoforms in long-read RNA-seq data [61–65]. However, analysis of single-nucleotide variants, such as RNA editing sites, presents significant challenges resulting from the relatively high level of sequencing errors. To this end, we present L-GIREMI, the first method, to our best knowledge, to identify RNA editing sites using long-read RNA-seq data.

L-GIREMI examines the linkage patterns between sequence variants in the same reads, complemented by a model-driven approach, to predict RNA editing sites. We adopted a similar strategy as in our previous method, GIREMI [66], which focused on short-read RNA-seq data. Long-read data is naturally advantageous in capturing correlative occurrence of multiple nucleotide variants in the mRNA. We showed that L-GIREMI affords high accuracy as reflected by the high fraction of A-to-G sites in its predictions. In addition, we demonstrated that the performance of L-GIREMI is robust given a wide range of total read coverage. Furthermore, as expected, RNA editing identification depends on the quality of the long-read data, with Sequel II-derived data out-performing those of the Sequel platform.

Long-read RNA-seq allows examination of co-occurrence of RNA editing sites in a single molecule. Leveraging this strength, we showed that editing sites in the same *Alu* or mRNA co-occurred more often than expected by chance. This observation extends previous reports using short-read RNA-seq that detected clustered RNA editing sites in hyper-edited regions [67]. Many scenarios may lead to co-occurrence of RNA editing sites, for instance long lifespan of RNA molecules, higher local concentration of ADAR proteins, or synergistic effects [68]. Nonetheless, it is important to note that this level of co-occurrence is lower than that driven by the linkage patterns of genetic variants on the same haplotype. Thus, the basic rationale of L-GIREMI that distinct MI distributions exist for genetic variants and RNA editing sites still holds.

Similar to RNA editing co-occurrence, long-read RNA-seq also allowed us to evaluate the possible existence of allele-specific editing. Our data supported the infrequent existence of this phenomenon. Allele-specific editing may be caused by SNP-related structural changes in the dsRNA region [69–71], demonstrating a mechanism of *cis*-regulation of RNA editing. The calculation of MI as implemented in L-GIREMI can be used to detect allele-specific editing events.

Another notable observation of our study is that dsRNA structure may lead to the skipping of a sizable region in the long-read RNA-seq. Previous studies also reported such observations, likely due to reverse transcriptase template switching, thus not unique to PacBio data [72–74]. This artifact may induce false positive calls of alternative splicing events, if the region-skipping is interpreted as a splicing event. It may also reduce the sensitivity in identifying RNA editing sites that reside in the skipped regions. On the other hand, compared to short-read data, long reads are uniquely advantageous in detecting this phenomenon, which may be leveraged to inform RNA secondary structure predictions in the future.

In summary, we present a method for the identification of RNA editing sites in long-read RNA-seq with high accuracy, even given low sequencing depth. Application of L-GIREMI allowed examination of RNA editing sites in single molecules, allele-specific RNA editing and region-skipping due to existence of dsRNA structures. This method provides a powerful means in examining nucleotide variants in long reads.

## Methods

### Mapping of reads using minimap2

Minimap2 was used for the mapping of long-read RNA-seq data against the human genome [44]. The ‘--cs’ option was included in order to output the cs tags that enabled parsing of sequence variants. Only unique mapped reads were retained for the identification of RNA editing sites. Samtools was used to remove multi-mapped reads with the option ‘-F 2052’ in the samtools view module [75]. Subsequently, the filtered SAM files were sorted according to genomic coordinates and converted into BAM files.

### The L-GIREMI analysis steps

The L-GIREMI algorithm consists of four main steps: 1. Correction of read orientation; 2. Collection of mismatches; 3. Calculation of mutual information among mismatch sites; 4. Scoring of mismatches with a generalized linear model (GLM). The details of each step is provided below. The source code of the program can be downloaded from https://github.com/gxiaolab/L-GIREMI.

#### Correction of read orientation

Although most reads generated by PacBio had the correct read strand, a minority of reads had wrong orientations, which can affect the accuracy of nucleotide variant analysis. We adopted the following strategy to check and, if necessary, correct the read strand. First, we obtained 3 types of strand information: the strand of the read from minimap2 (namely, mapped strand), the strand of the gene annotated (Gencode v34) at the read location (namely, annotated strand), and the strand from which the majority of the splice site sequences of the read were consistent with the known motifs (GT-AG, GC-AG or AT-AC) (namely, splice site strand). For most reads, all 3 sources gave the same strand. Otherwise, the read strand was set to be the dominant one among the three. Further, in rare cases, only two types of strand information was available due to the read mapping to intergenic regions or the lack of known splicing motif. For these cases, the annotated strand was given the highest priority, followed by splice site strand and mapped strand, respectively. If the read was mapped to a region with sense and antisense gene pairs, the gene with the larger overlap with the read sequence was used. It should be noted that reads harboring no spliced junctions were removed from the analysis to avoid contamination by DNA-derived reads.

#### Collection of mismatches

In this step, we obtain a catalog of all mismatches in the mapped reads, detected via the cs tags generated by minimap2. Their genomic coordinates, reference nucleotides, and alternative nucleotides were stored. In order to remove likely sequencing errors, several filters were implemented according to common practice in RNA editing identification via RNA-seq [76]. Specifically, the following mismatch sites were removed: (1) those with low read coverage (<3 reads). (2) Sites within simple repeats or homopolymers. (3) Sites within the vicinity (default 4 bp) of splice sites. (4) Sites with allelic ratio (minor allele/total) less than a threshold (default 0.1). Mismatched sites overlapping dbSNP annotations (v38) were labeled as SNPs and those with allelic ratio between a range (0.35 to 0.65 by default) were labeled as heterozygous SNPs, which were used in the calculation of mutual information.

#### Calculation of mutual information among mismatch sites

L-GIREMI calculates the mutual information (MI) for each mismatch site relative to heterozygous SNPs, and for pairs of heterozygous SNPs, respectively. For a mismatch (m_i_) and a SNP (s_j_), the MI (I) was calculated as:

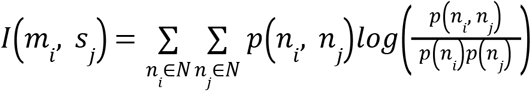

where *N* = {A, C, G, T}, and *n*_i_ and *n*_j_ denote the nucleotides of the sites in the pair. *p*(*n*_i_, *n*_j_) represents the probability of observing the *n*_i_ and *n*_j_ nucleotides in the same read, calculated using the maximum likelihood method. Natural logarithm was used for the calculation. The MI of a pair of SNPs was calculated similarly.

For every mismatch or SNP site, there might exist multiple other SNPs harbored within the same reads. Thus, the overall MI for the mismatch or SNP was calculated as the average of all the pairwise MI values. Empirical p values were calculated for all the mismatches based on the distribution of the SNP MI. Mismatches with p value smaller than a threshold (0.05 by default) and not included in the dbSNP database were selected as predicted RNA editing sites, which were used for GLM training described below.

#### Scoring of mismatches via a GLM

GLM was used as the scoring model of L-GIREMI. Features included in the model are the allelic ratio of the mismatch, the mismatch type and the sequences of the nearest neighbor nucleotides before and after the mismatch site. RNA editing sites identified in the above MI calculation step were used as positive training data. dbSNPs with p values larger than the threshold (0.05 by default) in the MI calculation step were used as negative training data. The training and testing of the GLM was carried out similarly as in GIREMI [77]. The score of each mismatch was calculated using the GLM. A score cutoff was chosen to maximize the F1 value. By default, predicted editing sites are defined as those that passed the score cutoff. This cutoff is dataset-specific. Alternatively, the user can define a customized score cutoff to achieve a desired level of %A-to-G among the predicted editing sites. As noted in the Results, for datasets with acceptable quality, the %A-to-G increased nearly monotonically relative to the GLM scores.

#### Calculation of the Gini index of Alu editing

The Gini index was calculated for each *Alu* using the editing ratio of each read. For each *Alu*, all possible editing sites were identified using all the reads that covered the *Alu*. Then, for each read, the fraction of possible editing sites that were edited in this read was calculated, referred to as the editing ratio of the read. The editing ratios of all reads for each *Alu* were then used to calculate the Gini index (G). In order to speed up the calculation, the Gini index was calculated as half of the mean absolute difference normalized by the mean of editing ratios [78]:

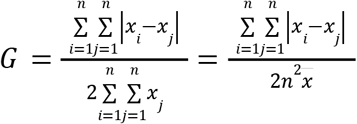

where *x*_i_ and *x*_j_ are the relative editing ratios of the read *i* and *j*, respectively, and *x* is the mean of all the editing ratios. *n* is the total number of reads for the *Alu*. This calculation was also carried out for shuffled data (based on randomization of As and Gs at possible editing sites across reads).

### Double-stranded RNA prediction

We applied a previously developed method to identify long dsRNAs harboring editing-enriched regions (EERs) [79]. This method is based on the rationale that the observation of A-to-I editing in an endogenous RNA is proof that the RNA is double stranded in vivo because ADARs edit dsRNAs. Briefly, we identified EERs using known RNA editing sites [80]. Overlapping 50bp windows (each with ≥3 editing sites) were combined and EERs within 2.5kb were classified as one EER. This distance allows formation of structures by distant binding partners. RNAfold was used to fold the EERs. Filters on minimum free energy and mismatch patterns were implemented to retain dsRNAs with >200bp stem length.

### Experimental validation of allele-specific editing via Sanger sequencing

Genomic DNA (gDNA) and total RNA were extracted from GM12878 cells using the Quick-DNA™ Miniprep kit (Zymo Research) and the Direct-zol™ RNA Miniprep Plus kit (Zymo Research), respectively, following the manufacturer’s protocols. 2ug of the total RNA was used to generate cDNA using the SuperScript™ IV First-Strand Synthesis System (Invitrogen). Sequences +/-100bp flanking the mismatch site were amplified using the DreamTaq PCR Master Mix (2X) (Thermo Scientific). The primers used for each amplicon (labeled as ZH1-7) are provided in Table S1. The PCR amplicons were resolved in 1% agarose gel and the bands of desired sizes were cut out and purified using the Zymoclean™ Gel DNA Recovery Kit (Zymo Research). Following purification, the amplicons were sent for Sanger sequencing (GENEWIZ from Azenta) using one of the PCR primers. Sites with alternative alleles in both gDNA and cDNA were validated as SNPs. Those with both A and G in the cDNA but only A in the gDNA were validated as RNA editing sites.

## Data and code availability

PacBio data derived from the brain sample of a patient with Alzheimer’s disease were downloaded from PacBio (https://downloads.pacbcloud.com/public/dataset/Alzheimer2019_IsoSeq/). PacBio data of GM12878 cells were downloaded from the ENCODE data portal (https://www.encodeproject.org/). L-GIREMI is available at https://github.com/gxiaolab/L-GIREMI

## Acknowledgement

We thank members of the Xiao laboratory for helpful discussions and comments on this work. This work was supported in part by grants U01HG009417 and R01MH123177 to XX, and UM1HG009443 to AM.

## Competing interests

The authors declare no competing interests.

**Figure S1**. Overview of the AD dataset. (**a**) Number of reads with mismatches, insertions, deletions or spliced junctions. The value next to the bar is the fraction of reads among all uniquely mapped reads. (**b**) Average number of mismatches, insertions or deletions per read.

**Figure S2**. Summary of mismatches observed in the AD dataset. (**a**) Mismatches obtained in the AD dataset after the pre-filtering step (step 2, Figure 1) in L-GIREMI. (**b**) %A-to-G among mismatches after the pre-filtering step in the subsets with different read coverages (via randomly subsampling of the AD dataset).

**Figure S3**. The data quality and RNA editing sites in the GM12878 long-read RNA-seq datasets generated by the Sequel II platform (ENCODE IDs: ENCFF417VHJ, ENCFF450VAU, ENCFF694DIE). (**a**) Number of reads with mismatches, insertions, deletions or spliced junctions. The value next to the bar is the fraction of reads among all uniquely mapped reads. (**b**) Average number of mismatches, insertions or deletions per read. (**c**) %A-to-G among all predicted editing sites vs. GLM scores. (**d**) RNA editing sites identified by L-GIREMI for the three datasets. The %A-to-G is shown in each graph. (**e**) %A-to-G among mismatches after the pre-filtering step (step 2, Figure 1) in L-GIREMI.

**Figure S4**. The data quality and RNA editing sites in the GM12878 long-read RNA-seq datasets generated by the Sequel platform (ENCODE IDs: ENCFF281TNJ, ENCFF475ORL, ENCFF329AYV, ENCFF902UIT). (**a-d**) Similar to Fig. S3 (a-d).

**Figure S5**. Cumulative distribution of mutual information of pairs of REDIportal editing sites or pairs of SNPs in the same gene. Compared to the shuffled controls, both editing sites and SNPs show higher levels of linkage (p < 0.001 for all comparisons, KS test) although the latter were associated with much higher mutual information.

**Figure S6**. Histogram of the MI for the editing sites identified in the ENCFF417VHJ dataset. Some sites had high MI (>0.3) as they were identified by the GLM step. Orange for sites in REDIportal, and gray for sites not in REDIportal.

**Figure S7**. Histograms of the read coverage of detected dsRNAs (n read ≥ 1) in the AD (top) or GM12878 data (bottom).

## References

1. Eisenberg E, Levanon EY. A-to-I RNA editing - immune protector and transcriptome diversifier. Nat Rev Genet. 2018;19:473–90.

2. Hough RF, Bass BL. Purification of the Xenopus laevis double-stranded RNA adenosine deaminase. J Biol Chem. Elsevier BV; 1994;269:9933–9.

3. Kim U, Garner TL, Sanford T, Speicher D, Murray JM, Nishikura K. Purification and characterization of double-stranded RNA adenosine deaminase from bovine nuclear extracts. J Biol Chem. Elsevier BV; 1994;269:13480–9.

4. Melcher T, Maas S, Herb A, Sprengel R, Seeburg PH, Higuchi M. A mammalian RNA editing enzyme. Nature. 1996;379:460–4.

5. O’Connell MA, Keller W. Purification and properties of double-stranded RNA-specific adenosine deaminase from calf thymus. Proc Natl Acad Sci U S A. 1994;91:10596–600.

6. Bass BL, Weintraub H. A developmentally regulated activity that unwinds RNA duplexes. Cell. Elsevier BV; 1987;48:607–13.

7. Nishikura K, Yoo C, Kim U, Murray JM, Estes PA, Cash FE, et al. Substrate specificity of the dsRNA unwinding/modifying activity. EMBO J. Wiley; 1991;10:3523–32.

8. Rebagliati MR, Melton DA. Antisense RNA injections in fertilized frog eggs reveal an RNA duplex unwinding activity. Cell. Elsevier BV; 1987;48:599–605.

9. Polson AG, Crain PF, Pomerantz SC, McCloskey JA, Bass BL. The mechanism of adenosine to inosine conversion by the double-stranded RNA unwinding/modifying activity: a high-performance liquid chromatography-mass spectrometry analysis. Biochemistry. 1991;30:11507–14.

10. Athanasiadis A, Rich A, Maas S. Widespread A-to-I RNA editing of Alu-containing mRNAs in the human transcriptome. PLoS Biol. 2004;2:e391.

11. Bazak L, Haviv A, Barak M, Jacob-Hirsch J, Deng P, Zhang R, et al. A-to-I RNA editing occurs at over a hundred million genomic sites, located in a majority of human genes. Genome Res. 2014;24:365–76.

12. Eisenberg E, Nemzer S, Kinar Y, Sorek R, Rechavi G, Levanon EY. Is abundant A-to-I RNA editing primate-specific? Trends Genet. 2005;21:77–81.

13. Kim DDY, Kim TTY, Walsh T, Kobayashi Y, Matise TC, Buyske S, et al. Widespread RNA editing of embedded alu elements in the human transcriptome. Genome Res. 2004;14:1719–25.

14. Jain M, Jantsch MF, Licht K. The Editor’s I on Disease Development. Trends Genet. 2019;35:903–13.

15. Li JB, Church GM. Deciphering the functions and regulation of brain-enriched A-to-I RNA editing. Nat Neurosci. 2013;16:1518–22.

16. Eisenberg E. Proteome Diversification by RNA Editing. Methods Mol Biol. 2021;2181:229–51.

17. Rueter SM, Dawson TR, Emeson RB. Regulation of alternative splicing by RNA editing. Nature. 1999;399:75–80.

18. Kapoor U, Licht K, Amman F, Jakobi T, Martin D, Dieterich C, et al. ADAR-deficiency perturbs the global splicing landscape in mouse tissues. Genome Res. 2020;30:1107–18.

19. Hsiao Y-HE, Bahn JH, Yang Y, Lin X, Tran S, Yang E-W, et al. RNA editing in nascent RNA affects pre-mRNA splicing. Genome Res. 2018;28:812–23.

20. Brümmer A, Yang Y, Chan TW, Xiao X. Structure-mediated modulation of mRNA abundance by A-to-I editing. Nat Commun. 2017;8:1255.

21. Stellos K, Gatsiou A, Stamatelopoulos K, Perisic Matic L, John D, Lunella FF, et al. Adenosine-to-inosine RNA editing controls cathepsin S expression in atherosclerosis by enabling HuR-mediated post-transcriptional regulation. Nat Med. 2016;22:1140–50.

22. Morita Y, Shibutani T, Nakanishi N, Nishikura K, Iwai S, Kuraoka I. Human endonuclease V is a ribonuclease specific for inosine-containing RNA. Nat Commun. 2013;4:2273.

23. Nishikura K. Editor meets silencer: crosstalk between RNA editing and RNA interference. Nat Rev Mol Cell Biol. 2006;7:919–31.

24. Wulff B-E, Nishikura K. Modulation of microRNA expression and function by ADARs. Curr Top Microbiol Immunol. 2012;353:91–109.

25. Liddicoat BJ, Piskol R, Chalk AM, Ramaswami G, Higuchi M, Hartner JC, et al. RNA editing by ADAR1 prevents MDA5 sensing of endogenous dsRNA as nonself. Science. 2015;349:1115–20.

26. Patterson JB, Samuel CE. Expression and regulation by interferon of a double-stranded-RNA-specific adenosine deaminase from human cells: evidence for two forms of the deaminase. Mol Cell Biol. 1995;15:5376–88.

27. Kim S, Ku Y, Ku J, Kim Y. Evidence of Aberrant Immune Response by Endogenous Double-Stranded RNAs: Attack from Within. Bioessays. 2019;41:e1900023.

28. Baysal BE, Sharma S, Hashemikhabir S, Janga SC. RNA Editing in Pathogenesis of Cancer. Cancer Res. 2017;77:3733–9.

29. Christofi T, Zaravinos A. RNA editing in the forefront of epitranscriptomics and human health. J Transl Med. 2019;17:319.

30. Krestel H, Meier JC. RNA Editing and Retrotransposons in Neurology. Front Mol Neurosci. 2018;11:163.

31. Bahn JH, Lee J-H, Li G, Greer C, Peng G, Xiao X. Accurate identification of A-to-I RNA editing in human by transcriptome sequencing. Genome Res. 2012;22:142–50.

32. Peng Z, Cheng Y, Tan BC-M, Kang L, Tian Z, Zhu Y, et al. Comprehensive analysis of RNA-Seq data reveals extensive RNA editing in a human transcriptome. Nat Biotechnol. 2012;30:253–60.

33. Ramaswami G, Lin W, Piskol R, Tan MH, Davis C, Li JB. Accurate identification of human Alu and non-Alu RNA editing sites. Nat Methods. 2012;9:579–81.

34. Park E, Williams B, Wold BJ, Mortazavi A. RNA editing in the human ENCODE RNA-seq data. Genome Res. 2012;22:1626–33.

35. Mansi L, Tangaro MA, Lo Giudice C, Flati T, Kopel E, Schaffer AA, et al. REDIportal: millions of novel A-to-I RNA editing events from thousands of RNAseq experiments. Nucleic Acids Res. 2021;49:D1012–9.

36. Zhang Q, Xiao X. Genome sequence-independent identification of RNA editing sites. Nat Methods. 2015;12:347–50.

37. Weirather JL, de Cesare M, Wang Y, Piazza P, Sebastiano V, Wang X-J, et al. Comprehensive comparison of Pacific Biosciences and Oxford Nanopore Technologies and their applications to transcriptome analysis. F1000Res. 2017;6:100.

38. Tardaguila M, de la Fuente L, Marti C, Pereira C, Pardo-Palacios FJ, Del Risco H, et al. SQANTI: extensive characterization of long-read transcript sequences for quality control in full-length transcriptome identification and quantification. Genome Res [Internet]. 2018; Available from: http://dx.doi.org/10.1101/gr.222976.117

39. Dong X, Tian L, Gouil Q, Kariyawasam H, Su S, De Paoli-Iseppi R, et al. The long and the short of it: unlocking nanopore long-read RNA sequencing data with short-read differential expression analysis tools. NAR Genom Bioinform. 2021;3:lqab028.

40. Tang AD, Soulette CM, van Baren MJ, Hart K, Hrabeta-Robinson E, Wu CJ, et al. Full-length transcript characterization of SF3B1 mutation in chronic lymphocytic leukemia reveals downregulation of retained introns. Nat Commun. 2020;11:1438.

41. Tardaguila M, de la Fuente L, Marti C, Pereira C, Pardo-Palacios FJ, Del Risco H, et al. SQANTI: extensive characterization of long-read transcript sequences for quality control in full-length transcriptome identification and quantification. Genome Res [Internet]. 2018; Available from: http://dx.doi.org/10.1101/gr.222976.117

42. Wyman D, Balderrama-Gutierrez G, Reese F, Jiang S, Rahmanian S, Forner S, et al. A technology-agnostic long-read analysis pipeline for transcriptome discovery and quantification [Internet]. bioRxiv. bioRxiv; 2019. Available from: http://biorxiv.org/lookup/doi/10.1101/672931

43. Zhang Q, Xiao X. Genome sequence-independent identification of RNA editing sites. Nat Methods. 2015;12:347–50.

44. Li H. Minimap2: pairwise alignment for nucleotide sequences. Bioinformatics. 2018;34:3094–100.

45. Lee J-H, Ang JK, Xiao X. Analysis and design of RNA sequencing experiments for identifying RNA editing and other single-nucleotide variants. RNA. 2013;19:725–32.

46. Quinones-Valdez G, Tran SS, Jun H-I, Bahn JH, Yang E-W, Zhan L, et al. Regulation of RNA editing by RNA-binding proteins in human cells. Commun Biol. 2019;2:19.

47. Weirather JL, de Cesare M, Wang Y, Piazza P, Sebastiano V, Wang X-J, et al. Comprehensive comparison of Pacific Biosciences and Oxford Nanopore Technologies and their applications to transcriptome analysis. F1000Res. 2017;6:100.

48. Ramaswami G, Lin W, Piskol R, Tan MH, Davis C, Li JB. Accurate identification of human Alu and non-Alu RNA editing sites. Nat Methods. 2012;9:579–81.

49. Ramaswami G, Zhang R, Piskol R, Keegan LP, Deng P, O’Connell MA, et al. Identifying RNA editing sites using RNA sequencing data alone. Nat Methods. 2013;10:128–32.

50. Roth SH, Levanon EY, Eisenberg E. Genome-wide quantification of ADAR adenosine-to-inosine RNA editing activity. Nat Methods. 2019;16:1131–8.

51. Weirather JL, de Cesare M, Wang Y, Piazza P, Sebastiano V, Wang X-J, et al. Comprehensive comparison of Pacific Biosciences and Oxford Nanopore Technologies and their applications to transcriptome analysis. F1000Res. 2017;6:100.

52. Bahn JH, Lee J-H, Li G, Greer C, Peng G, Xiao X. Accurate identification of A-to-I RNA editing in human by transcriptome sequencing. Genome Res. 2012;22:142–50.

53. Zhou Z-Y, Hu Y, Li A, Li Y-J, Zhao H, Wang S-Q, et al. Genome wide analyses uncover allele-specific RNA editing in human and mouse. Nucleic Acids Res. 2018;46:8888–97.

54. Cocquet J, Chong A, Zhang G, Veitia RA. Reverse transcriptase template switching and false alternative transcripts. Genomics. 2006;88:127–31.

55. Blango MG, Bass BL. Identification of the long, edited dsRNAome of LPS-stimulated immune cells. Genome Res. 2016;26:852–62.

56. Holmqvist I, Bäckerholm A, Tian Y, Xie G, Thorell K, Tang K-W. FLAME: long-read bioinformatics tool for comprehensive spliceome characterization. RNA. 2021;27:1127–39.

57. Tang AD, Soulette CM, van Baren MJ, Hart K, Hrabeta-Robinson E, Wu CJ, et al. Full-length transcript characterization of SF3B1 mutation in chronic lymphocytic leukemia reveals downregulation of retained introns. Nat Commun. 2020;11:1438.

58. Tardaguila M, de la Fuente L, Marti C, Pereira C, Pardo-Palacios FJ, Del Risco H, et al. SQANTI: extensive characterization of long-read transcript sequences for quality control in full-length transcriptome identification and quantification. Genome Res [Internet]. 2018; Available from: http://dx.doi.org/10.1101/gr.222976.117

59. Tian L, Jabbari JS, Thijssen R, Gouil Q, Amarasinghe SL, Kariyawasam H, et al. Comprehensive characterization of single cell full-length isoforms in human and mouse with long-read sequencing [Internet]. bioRxiv. 2020 [cited 2022 Feb 26]. p. 2020.08.10.243543. Available from: https://www.biorxiv.org/content/10.1101/2020.08.10.243543v1.abstract

60. Wyman D, Balderrama-Gutierrez G, Reese F, Jiang S, Rahmanian S, Forner S, et al. A technology-agnostic long-read analysis pipeline for transcriptome discovery and quantification [Internet]. bioRxiv. bioRxiv; 2019. Available from: http://biorxiv.org/lookup/doi/10.1101/672931

61. Holmqvist I, Bäckerholm A, Tian Y, Xie G, Thorell K, Tang K-W. FLAME: long-read bioinformatics tool for comprehensive spliceome characterization. RNA. 2021;27:1127–39.

62. Tang AD, Soulette CM, van Baren MJ, Hart K, Hrabeta-Robinson E, Wu CJ, et al. Full-length transcript characterization of SF3B1 mutation in chronic lymphocytic leukemia reveals downregulation of retained introns. Nat Commun. 2020;11:1438.

63. Tardaguila M, de la Fuente L, Marti C, Pereira C, Pardo-Palacios FJ, Del Risco H, et al. SQANTI: extensive characterization of long-read transcript sequences for quality control in full-length transcriptome identification and quantification. Genome Res [Internet]. 2018; Available from: http://dx.doi.org/10.1101/gr.222976.117

64. Tian L, Jabbari JS, Thijssen R, Gouil Q, Amarasinghe SL, Kariyawasam H, et al. Comprehensive characterization of single cell full-length isoforms in human and mouse with long-read sequencing [Internet]. bioRxiv. 2020 [cited 2022 Feb 26]. p. 2020.08.10.243543. Available from: https://www.biorxiv.org/content/10.1101/2020.08.10.243543v1.abstract

65. Wyman D, Balderrama-Gutierrez G, Reese F, Jiang S, Rahmanian S, Forner S, et al. A technology-agnostic long-read analysis pipeline for transcriptome discovery and quantification [Internet]. bioRxiv. bioRxiv; 2019. Available from: http://biorxiv.org/lookup/doi/10.1101/672931

66. Zhang Q, Xiao X. Genome sequence-independent identification of RNA editing sites. Nat Methods. 2015;12:347–50.

67. Porath HT, Carmi S, Levanon EY. A genome-wide map of hyper-edited RNA reveals numerous new sites. Nat Commun. 2014;5:4726.

68. Rodriques SG, Chen LM, Liu S, Zhong ED, Scherrer JR, Boyden ES, et al. RNA timestamps identify the age of single molecules in RNA sequencing. Nat Biotechnol. 2021;39:320–5.

69. Cocquet J, Chong A, Zhang G, Veitia RA. Reverse transcriptase template switching and false alternative transcripts. Genomics. 2006;88:127–31.

70. Houseley J, Tollervey D. Apparent non-canonical trans-splicing is generated by reverse transcriptase in vitro. PLoS One. 2010;5:e12271.

71. Tardaguila M, de la Fuente L, Marti C, Pereira C, Pardo-Palacios FJ, Del Risco H, et al. SQANTI: extensive characterization of long-read transcript sequences for quality control in full-length transcriptome identification and quantification. Genome Res [Internet]. 2018; Available from: http://dx.doi.org/10.1101/gr.222976.117

72. Cocquet J, Chong A, Zhang G, Veitia RA. Reverse transcriptase template switching and false alternative transcripts. Genomics. 2006;88:127–31.

73. Houseley J, Tollervey D. Apparent non-canonical trans-splicing is generated by reverse transcriptase in vitro. PLoS One. 2010;5:e12271.

74. Tardaguila M, de la Fuente L, Marti C, Pereira C, Pardo-Palacios FJ, Del Risco H, et al. SQANTI: extensive characterization of long-read transcript sequences for quality control in full-length transcriptome identification and quantification. Genome Res [Internet]. 2018; Available from: http://dx.doi.org/10.1101/gr.222976.117

75. Li H, Handsaker B, Wysoker A, Fennell T, Ruan J, Homer N, et al. The Sequence Alignment/Map format and SAMtools. Bioinformatics. 2009;25:2078–9.

76. Quinones-Valdez G, Tran SS, Jun H-I, Bahn JH, Yang E-W, Zhan L, et al. Regulation of RNA editing by RNA-binding proteins in human cells. Commun Biol. 2019;2:19.

77. Zhang Q, Xiao X. Genome sequence-independent identification of RNA editing sites. Nat Methods. 2015;12:347–50.

78. Sen A, Sen A, Sen MA, Foster JE, Amartya S,. Foster JE, et al. On Economic Inequality. Oxford University Press; 1997.

79. Sen A, Sen A, Sen MA, Foster JE, Amartya S,. Foster JE, et al. On Economic Inequality. Oxford University Press; 1997.

80. Lo Giudice C, Tangaro MA, Pesole G, Picardi E. Investigating RNA editing in deep transcriptome datasets with REDItools and REDIportal. Nat Protoc. 2020;15:1098–131.

