## Supplementary Figures for "L-GIREMI uncovers RNA editing sites in long-read RNA-seq"

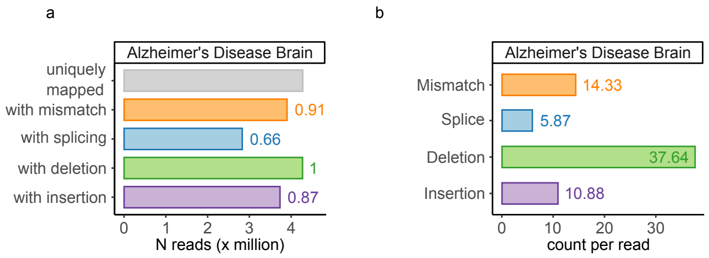


Figure S1: Overview of the AD dataset. (a) Number of reads with mismatches, insertions, deletions or spliced junctions. The value next to the bar is the fraction of reads among all uniquely mapped reads. (b) Average number of mismatches, insertions or deletions per read.

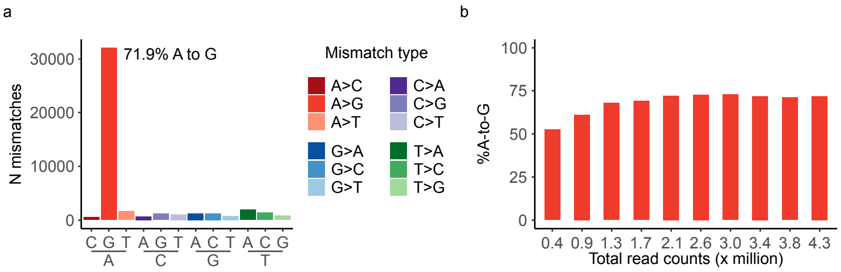


Figure S2: Summary of mismatches observed in the AD dataset. (a) Mismatches obtained in the AD dataset after the pre-filtering step (step 2, Figure 1) in L-GIREMI. (b) %A-to-G among mismatches after the pre-filtering step in the subsets with different read coverages (via randomly subsampling of the AD dataset).

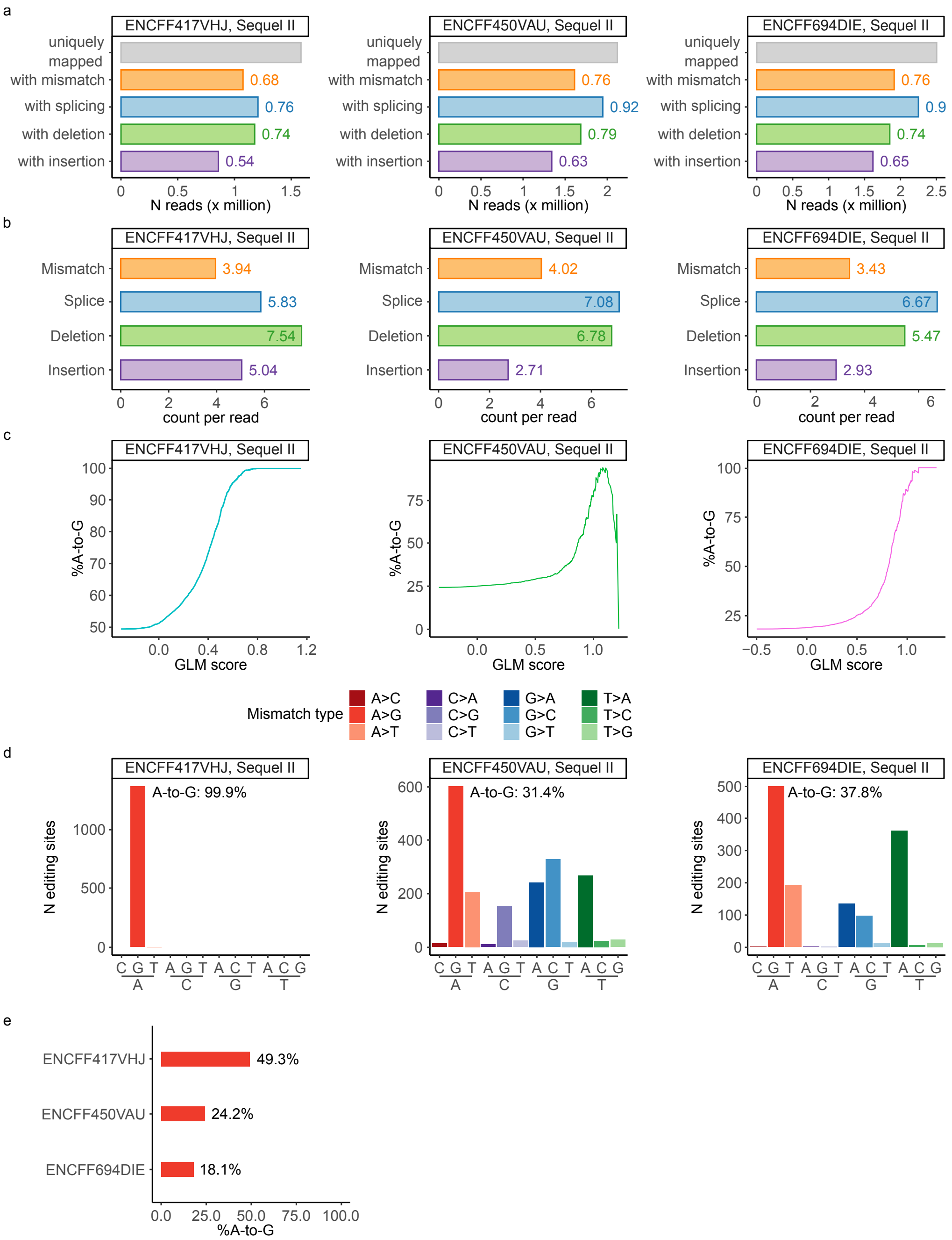


Figure S3: The data quality and RNA editing sites in the GM12878 long-read RNA-seq datasets generated by the Sequel II platform (ENCODE IDs: ENCFF417VHJ, ENCFF450VAU, ENCFF694DIE). (a) Number of reads with mismatches, insertions, deletions or spliced junctions. The value next to the bar is the fraction of reads among all uniquely mapped reads. (b) Average number of mismatches, insertions or deletions per read. (c) %A-to-G among all predicted editing sites vs. GLM scores. (d) RNA editing sites identified by L-GIREMI for the three datasets. The %A-to-G is shown in each graph. (e) %A-to-G among mismatches after the pre-filtering step (step 2, Figure 1) in L-GIREMI.

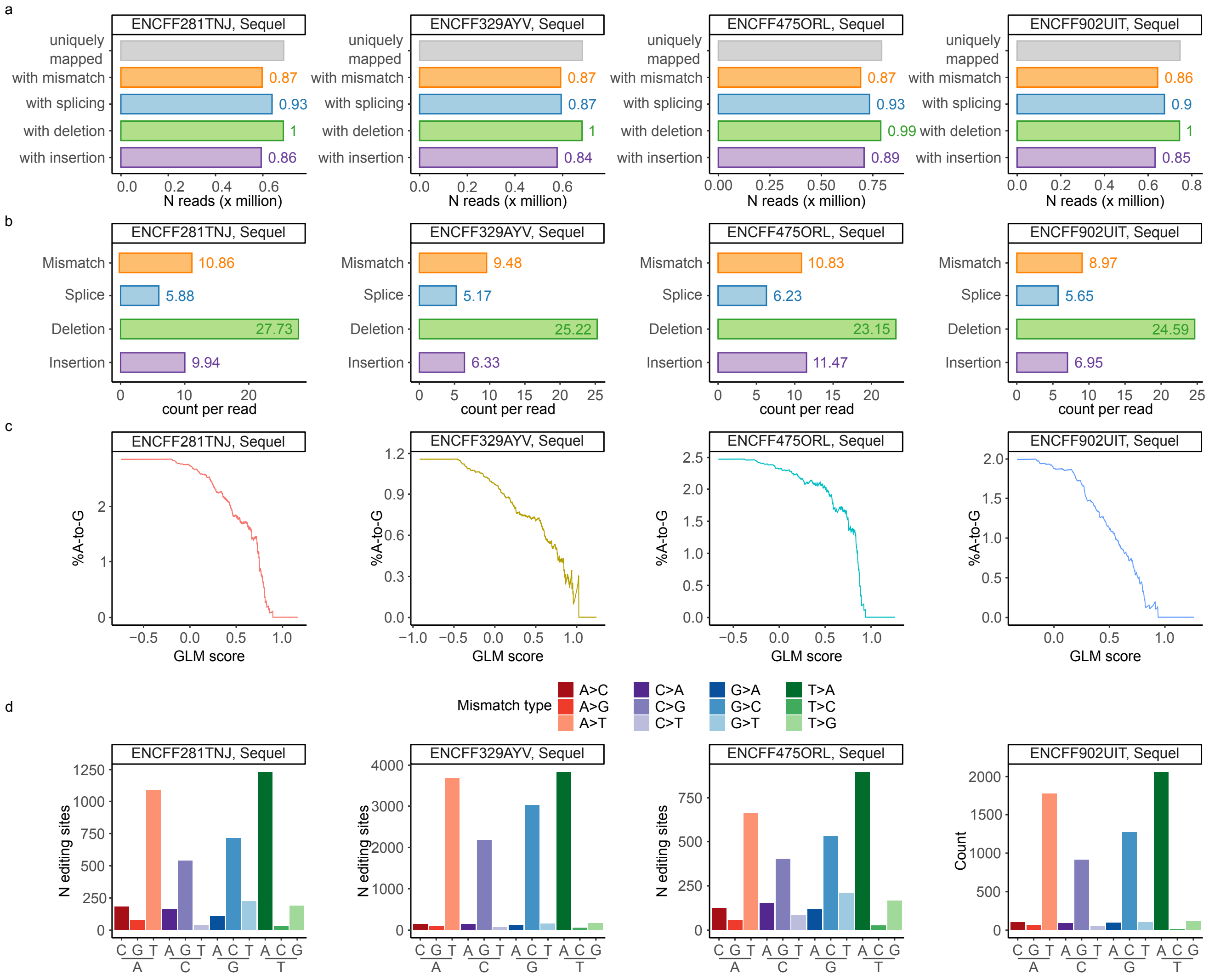


Figure S4: The data quality and RNA editing sites in the GM12878 long-read RNA-seq datasets generated by the Sequel platform (ENCODE IDs: ENCFF281TNJ, ENCFF475ORL, ENCFF329AYV, ENCFF902UIT). (a-d) Similar to Fig. S3 (a-d).

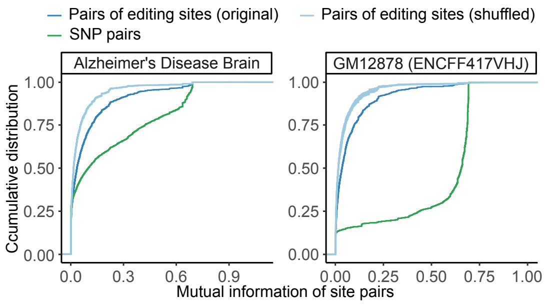


Figure S5: Cumulative distribution of mutual information of pairs of REDIportal editing sites or pairs of SNPs in the same gene. Compared to the shuffled controls, both editing sites and SNPs show higher levels of linkage ($p < 0.001$ for all comparisons, KS test) although the latter were associated with much higher mutual information.

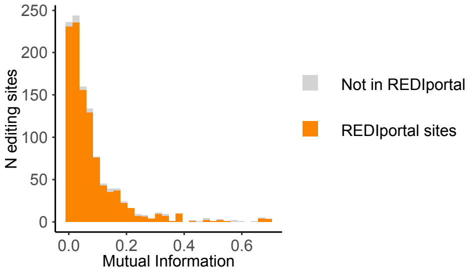


Figure S6: Histogram of the MI for the editing sites identified in the ENCFF417VHJ dataset. Orange for sites in REDIportal, and gray for sites not in REDIportal.

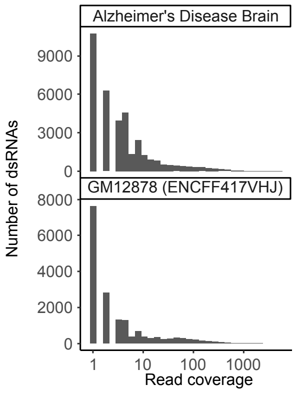


Figure S7: Histograms of the read coverage of detected dsRNAs ($nread \leq 1$) in the AD (top) or GM12878 data (bottom).
